# Exploration of the sensitivity to macrocyclic lactones in the canine heartworm (*Dirofilaria immitis*) in Australia using phenotypic and genotypic approaches

**DOI:** 10.1101/2022.10.15.512343

**Authors:** Rosemonde Isabella Power, Jan Šlapeta

## Abstract

Canine heartworm disease is a potentially deadly cardiopulmonary disease caused by the mosquito-borne filarial nematode *Dirofilaria immitis*. In Australia, the administration of macrocyclic lactone (ML) drugs has successfully reduced the prevalence of *D. immitis* infection. However, the recent re-emergence of *D. immitis* in dogs in Queensland, Australia and the identification of ML-resistant isolates in the USA poses an important question of whether ML-resistance has emerged in this parasite in Australia. The aim of this study was to utilise phenotypic and genotypic approaches to examine the sensitivity to ML drugs in *D. immitis* in Australia. To do this, we surveyed 45 dogs from Queensland and New South Wales across 3 years (2019-2022) for the presence of *D. immitis* infection using an antigen test, quantitative Modified Knott’s test, and qPCR targeting both *D. immitis* and the *D. immitis* symbiont *Wolbachia*. A phenotype observed by utilising sequential quantification of microfilariae for 23/45 dogs was coupled with genetic testing of filtered microfilariae for SNPs previously associated with ML-resistance in isolates from the USA. Sixteen (16/45) dogs tested positive for *D. immitis* infection despite reportedly receiving ‘rigorous’ heartworm prevention for 12 months prior to the study, according to the owners’ assessment. The phenotype and genotypic assays in this study did not unequivocally demonstrate the presence of ML-resistant *D. immitis* in Australia. Although the failure of 16 dogs to reduce microfilaremia by >90% after ML treatment was considered a suspect phenotype of ML-resistance, no genotypic evidence was discovered using the genetic SNP analysis. The traditional quantitative Modified Knott’s test can be substituted by qPCR targeting *D. immitis* or associated *Wolbachia* endosymbiont DNA for a more rapid measurement of microfilariae levels. More definitive phenotypic evidence of resistance is critically needed before the usefulness of SNPs for the detection of ML-resistance in Australia can be properly assessed.

## 1. Introduction

Canine heartworm disease is a potentially deadly cardiopulmonary disease caused by the mosquito-borne filarial nematode *Dirofilaria immitis*, which is distributed across many parts of the world (Simón et al., 2012; Genchi and Kramer, 2020; Lau et al., 2021; Noack et al., 2021). Although the prevalence of *D. immitis* in Australia is low, the parasite is endemic to the country and has recently re-emerged in dogs in Queensland (Nguyen et al., 2016; Orr et al., 2020; Panetta et al., 2021). For the past 30 years in Australia and the USA, the main strategy for canine heartworm prevention has been the administration of macrocyclic lactone (ML) anthelmintics (Orr et al., 2020; Diakou and Prichard, 2021; Lau et al., 2021; Noack et al., 2021; Prichard, 2021). These ML preventatives include monthly oral or topical formulations of ivermectin, milbemycin oxime, moxidectin and selamectin, or extended-release injections of moxidectin microspheres (Nguyen et al., 2016; McTier et al., 2019a). When used according to product instructions, these preventatives are approved to be 100% effective against less than 30-day old *D. immitis* (Hampshire, 2005). Over the past 20 years, lack of efficacy (LOE) reports for all heartworm product categories have occurred in the Lower Mississippi River Valley region of the USA (Hampshire, 2005). Most of these LOE reports were associated with compliance issues and extenuating circumstances, which prevented dogs from receiving sufficient product every 30 days (Atkins et al., 2014). A small number of LOE cases were not linked to noncompliance, indicating the potential development of ML-resistance in canine heartworms (Atkins et al., 2014). Laboratory studies have now identified several ML-resistant *D. immitis* isolates circulating in the Lower Mississippi Valley region of the USA (Blagburn et al., 2011; Pulaski et al., 2014; Bourguinat et al., 2015; Blagburn et al., 2016; Maclean et al., 2017; McTier et al., 2017b; Diakou and Prichard, 2021). These studies involved experimentally infecting ML-naïve dogs with infective L3 obtained from field LOE cases (Pulaski et al., 2014). In addition, <100% efficacy has been experimentally demonstrated for all currently marketed MLs (Bourguinat et al., 2015; Blagburn et al., 2016; McTier et al., 2019b).

MLs are essentially the only class of drugs currently available for canine heartworm prevention in Australia, in addition to the daily diethylcarbamazine citrate (Noack et al., 2021). Hence, the emergence of ML-resistance in *D. immitis* in the USA presents a significant concern for the veterinary industry by limiting the usefulness of ML products and jeopardising canine health. Prolonging the efficacy of MLs for canine heartworm prevention is warranted. With the absence of rapid biological tests that would demonstrate LOE of the target population (<30-day old *D. immitis*), discovery of heritable DNA molecular markers is needed for monitoring ML-resistance in adult *D. immitis* populations or their microfilariae within the dog bloodstream. Previous studies from the USA have compared the genotypes of different groups of *D. immitis* and identified single nucleotide polymorphisms (SNPs) potentially linked to ML-resistance (Bourguinat et al., 2011; Bourguinat et al., 2015). A whole genome approach was used in Bourguinat et al. (2015) to identify 186 SNP loci associated with the inability of dogs to remove circulating microfilariae after ML administration. From these 186 SNPs, a subset of 42 SNPs was initially validated as being highly associated with the ML-resistance phenotype in *D. immitis* (Bourguinat et al., 2015). The 42 SNPs were further evaluated in 10 additional ML-susceptible and ML-resistant isolates to generate 2, 3, 5 and 10-SNP predictive models for ML-resistance (Bourguinat et al., 2017). The 10 SNP approach that most effectively separated ML-susceptible and ML-resistant isolates was validated in an extensive study from south-east USA with field-collected *D. immitis* microfilariae, demonstrating high levels of sensitivity and specificity (Ballesteros et al., 2018). The SNP-based genotype predicted the phenotype which was defined by the inability of MLs to clear microfilariae using a suppression test and thus implied the presence of ML-resistant *D. immitis* infection (Ballesteros et al., 2018).

In contrast to the USA, it is currently unknown whether ML-resistance has emerged in *D. immitis* in Australia. A recent study revealed a large proportion of Australian dogs that were microfilaria-positive despite ‘rigorous’ administration of MLs in far-north Queensland (Nguyen et al., 2016). However, evidence of ML purchase history or ML administration was not provided by clients (Nguyen et al., 2016). A study that adopted the predictive SNP models developed in the USA to determine ML-resistance status did not find evidence of SNPs associated with ML-resistance in adult *D. immitis* from Sydney, Australia (Lau et al., 2021). As *D. immitis* prevention remains critical and prevalent across tropical Australia, further inquiry into ML-resistance in Australian *D. immitis* populations is necessary (Orr et al., 2020; Panetta et al., 2021). New screening tools and surveillance are required that apply or adapt the microfilarial suppression test originally proposed by Geary et al. (2011) and subsequently utilised by Ballesteros et al. (2018).

The aim of this study was to utilise phenotypic and genotypic approaches to determine the presence of ML-resistant *D. immitis* in Australia. To do so, we surveyed dogs across 3 years (2019-2022) for the presence of *D. immitis* using an antigen test, quantitative Modified Knott’s test, and qPCR targeting both *D. immitis* as well as the *D. immitis* symbiont *Wolbachia*. Material was collected prior to ML application and post-ML application at varying intervals. A Modified Knott’s test was coupled with filtration of microfilariae for genetic testing of SNPs previously associated with ML-resistance in isolates from the USA.

## 2. Material & Methods

### 2.1. Blood collection from dogs in Queensland and New South Wales

Blood samples were collected opportunistically from dogs in Queensland and New South Wales, Australia between September 2019 and January 2022. The dogs selected for this study were privately-owned animals from participating veterinary practices where the practitioner had either confirmed or suspected the presence of *D. immitis* infections. Blood samples (≤4 mL) were collected by registered veterinarians and shipped in EDTA tubes to the Sydney School of Veterinary Science, University of Sydney, Australia for diagnostic *D. immitis* testing. The following details were obtained for each dog: sex, age, breed, desexing status, locality, heartworm prevention history for the previous 12 months based on the owner’s assessment, and whether the animal had lived or visited other states or territories. Dog owners were required to complete a consent form prior to blood collection.

### 2.2. Sequential Modified Knott’s test

To determine the effect of ML administration on *D. immitis* microfilariae counts from infected dogs, we utilised a Modified Knott’s test. This was used to enumerate the number of *D. immitis* microfilariae in dog blood before ML treatment and then at a second point following the ML treatment. The time interval between drug treatment and the follow-up count varied across the study as necessitated by the need to follow timelines within the veterinary practices that were providing blood samples for the study, but was generally in the period 30-40 days after treatment. In some cases, an additional post-treatment microfilariae count was made at a later timepoint. The type of ML product administered was at the veterinarian’s discretion, according to their best practice in heartworm disease management. We calculated the percentage (%) reduction of microfilariae between the pre-treatment and post-treatment samples. In cases in which dogs showed <90% reductions in microfilariae, additional blood samples were collected where possible to observe whether microfilariae levels eventually declined.

To perform the Modified Knott’s test, 1 mL of whole blood was mixed with 10 mL of 2% formalin in a 15-mL tube (Knott, 1939; Mylonakis et al., 2004). The mixture was centrifuged at 2,300 relative centrifugal force (RCF) for 5 min using a benchtop centrifuge (POCD Scientific, Australia). The pellet was mixed with 1% methylene blue stain and enumerated microscopically. The slide was examined under a microscope (BX40, Olympus, Australia) using 10× objective and the number of microfilariae observed in the entire slide was counted.

### 2.3. Detection of *D. immitis* antigen using DiroCHEK^®^ ELISA

Prior to performing DiroCHEK^®^ ELISA (Zoetis, Australia) testing, the stored plasma samples were thawed at room temperature and vortexed for 3 sec. Both heated and unheated (neat) aliquots of each plasma sample were tested. To heat the plasma samples, a 100 μL aliquot of plasma was diluted with 100 μL of phosphate buffered saline (PBS; pH = 7.4) in a plain 1.5-mL tube. For samples with insufficient plasma volume, only 50 μL of plasma was diluted with 50 μL of PBS. The diluted plasma samples were heated at 103°C for 10 min using a dry heat block (Major Science). Heated samples were immediately centrifuged at 16,100 RCF for 5 min using a benchtop centrifuge (Eppendorf, Australia) set at 4°C. The supernatant was transferred into a separate 1.5-mL tube for use in DiroCHEK^®^ ELISA testing. To load the DiroCHEK^®^ ELISA plate, 50 μL of heated and unheated plasma for each sample were transferred into separate wells. Positive and negative controls provided by the manufacturer were included, and samples with leftover heated plasma were included as repeats. The DiroCHEK^®^ ELISA test was then performed according to manufacturer’s instructions. The optical density (OD) of each sample in the DiroCHEK^®^ ELISA plate was measured at 620 nm in a Halo LED 96 microplate reader (Dynamica). To distinguish between negative and positive samples, the following equation was applied as a threshold:

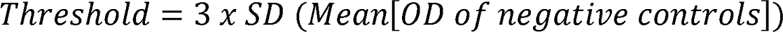

### 2.4. Microfilariae extraction

To isolate microfilariae from dog blood, a filtration procedure was adopted and modified (Bourguinat et al., 2015). Prior to filtration, 1 mL of whole blood was diluted with 35 mL of Red Blood Cell lysis buffer (8 g/L sodium carbonate, 5 mL/L Triton X-100) in a 50-mL tube. For one sample (HW52), only 100 μl of blood was diluted. The diluted blood was then pushed through a syringe containing a polycarbonate membrane filter (pore size 20 µm, filter diameter 25 mm; Sterlitech^®^ Corporation, WA). This allowed the microfilariae to be collected on the filter membrane whilst other blood components passed through the syringe. The syringe was then flushed with 50 mL of phosphate buffered saline (PBS). The filter membrane was removed from the syringe, placed into a 1.5-mL tube, and stored at -20°C until further use.

### 2.5. DNA isolation

DNA was isolated from 100 µL of whole blood using the blood protocol from the Monarch^®^ Genomic DNA Purification Kit (New England Biolabs, Australia). In addition, DNA was isolated from microfilariae on filter membranes using the same kit. DNA isolation from filter membranes followed the animal tissue sample protocol and used Proteinase K. Each DNA isolation batch contained an extraction blank. The eluted DNA was stored at -20°C until further use in the qPCR and next generation sequencing (NGS) assays.

### 2.6. Quantitative PCR targeting dog, *D. immitis* and *Wolbachia* sp. of *D. immitis*

A qPCR targeting the partial canine glyceraldehyde3-phosphate dehydrogenase (*GAPDH*) region was performed to verify the presence of dog DNA using primers with a probe, as previously described (Orr et al., 2020; Panetta et al., 2021) (Supplementary Table S1). A diagnostic TaqMan probe qPCR targeting the *cox1* gene was performed to detect *D. immitis* DNA using primers with a probe, as previously described (Panetta et al., 2021) (Supplementary Table S1). A qPCR targeting the *ftsZ* gene of *D. immitis*-associated *Wolbachia* was performed using primers and probe described in Laidoudi et al. (2020) (Supplementary Table S1).

The dog, *D. immitis* and *Wolbachia* qPCR assays were carried out in 10 μL final volumes of Luna® Universal Probe qPCR Master Mix (New England Biolabs, Australia) containing 1 μL of template DNA isolated from whole dog blood and primers and probe at a final concentration of 400 nM and 100 nM, respectively. To confirm the presence of *D. immitis*, qPCRs targeting dog and *D. immitis* DNA were repeated using DNA from extracted microfilariae. For these reactions, the volumes of primer, probe, PCR-grade water, and template DNA were doubled to reach final volumes of 20 μL. All qPCRs were carried out on a CFX96 Real-Time PCR system with the corresponding Bio-Rad CFX Maestro Software (BioRad, Australia). Cycling conditions for all three assays consisted of an initial denaturation at 95°C for 1 min followed by 40 two-step cycles at 95°C for 15 sec and 60°C for 30 sec. Each qPCR run included DNA extraction blanks and a no template control (NTC) of PCR-grade water. The arbitrary qPCR baseline threshold was set to 100 RFU and C_t_-values were reported for all samples. A qPCR targeting the *GAPDH* region of the dog genome confirmed the presence of dog DNA in all samples. All No Template Control (NTC) reactions remained negative.

### 2.7. Statistical analysis of phenotypic data

A paired t-test was performed to compare the OD between neat and heated plasma samples in the DiroCHEK^®^ ELISA. Additional paired t-tests were performed to observe differences between pre- and post-ML treatment samples in the qPCRs. To evaluate linear relationships between phenotypic datasets, Pearson’s correlation coefficients were generated. Prior to performing correlation analyses with the qPCR data, ‘N/A’ results were replaced with a C_t_-value of ‘40’. All statistical analyses were performed using GraphPad Prism (version 9.3.1).

### 2.8. Selection of SNP markers

Studies from the USA identified 10 SNPs that best differentiated ML-susceptible *D. immitis* from ML-resistant isolates (Bourguinat et al., 2017). Seven of these SNPs were selected for analysis in this study as they covered both 5-SNP models generated by Ballesteros et al. (2018) and Bourguinat et al. (2017). The 2 SNP markers (L15709_A, L30575) reported as giving the best prediction of ML-resistance in Ballesteros et al. (2018) were included in these 7 SNPs. Additional information about the 7 SNPs selected for this study including their position in the genome and various names used in other studies are available in Supplementary Table S2.

### 2.9. Illumina PCR amplicon Next Generation Sequencing targeting 7 SNPs associated with ML-resistance in USA

Illumina overhang adapters were added to all 7 primer sets which targeted regions encompassing the SNPs of interest (L42411, L21554, L45689, L9400, L20587, L15709_A, and L30575) according to Bourguinat et al. (2015). Their reference sequences were mapped to *D. immitis* genome nDi.2.2 scaffolds (Godel et al., 2012). The first stage PCRs for Illumina Next Generation Sequencing (NGS) were performed in-house according to the ‘16S Metagenomic Sequencing Library Preparation’ instructions for the Illumina MiSeq system (Illumina, Australia). Conventional PCR assays were carried out with Q5 Hot Start High-Fidelity 2× Master Mix (New England Biolabs, Australia). For some samples, dilutions of 1:10 or 1:100 were used due to low C_t_-values obtained from the *D. immitis*-specific qPCR. Conventional PCRs were performed on a Bio-Rad T100™ Thermal Cycler (Bio-Rad, Australia). The cycling conditions included an initial denaturation step at 98°C for 30 sec followed by 27 – 35 cycles of 98°C for 5 sec, 55°C for 10 sec and 72°C for 20 sec with a final extension at 72°C for 2 min. The number of cycles varied so that each sample reached the exponential phase of amplification, as previously described (Power et al., 2021). To increase efficiency of the first stage PCRs, conventional PCR assays were replaced with qPCR assays with SensiFAST™ SYBR® No-ROX Kit (Bioline, Australia). The cycling conditions included an initial denaturation at 95°C for 3 min followed by 28 -40 cycles of 95°C for 5 sec, 55°C for 10 sec and 72°C for 20 sec with a final melt curve. The inclusion of a melt curve allowed us to confirm the presence of the desired PCR product. DNA of an adult *D. immitis* collected from a dog in Sydney (P6/20D), Australia was used as a positive control (Lau et al., 2021). A no template control (NTC) of PCR-grade water was included for each SNP assay. Presence of expected PCR products was confirmed by visualisation on agarose gel for conventional PCR and qPCR products with ambiguous melt profiles. To determine whether comparable results were obtained using the two PCR approaches, three samples (HW19, HW20, HW32) were run as both conventional PCR and qPCR reactions for 5 loci (L42411, 21554, 45689, 9400, 20587), along with additional samples for L42411 (HW33) and L20587 (HW63). To investigate the impact of DNA concentration on the NGS results, four samples in the L45689 assay (HW20, HW41, HW45, HW66) and four samples in the L9400 assay (HW20, HW31, HW66, P6/20D) were repeated using alternative dilutions or cycle numbers. As one sample (HW52) was analysed using filtered microfilariae from 100 µL of blood, repeats for three loci (L20587, L15709A, L30575) were included using filtered microfilariae from 1 mL of blood. There was no difference that wound compromised SNP calls between the approaches irrespective of the dilution, blood volume, cycle number or PCR approach used.

After the first-stage PCR, products were submitted to the Ramaciotti Centre for Genomics, University of New South Wales, Australia for indexing (384), library preparation and next generation sequencing (NGS) using Illumina MiSeq v2 250PE sequencing run.

### 2.10. SNP analysis of NGS samples

Data as FastQ files were analysed in R 4.0.5 within RStudio 2022.07.1 Build 554 (RStudio Team, 2022). For each SNP locus, primers were identified and removed from all sequencing reads using Cutadapt 4.0 (Martin, 2011) followed by DADA2, an open-source R package (https://github.com/benjjneb/dada2). DADA2 pipeline (1.18.0; Callahan et al. (2016)) was used to infer the *D. immitis* SNP amplicon sequence variants (ASVs) present in the filtered microfilariae DNA samples. The quality profiles of the cut sequencing reads were visualised and inspected using the “plotQualityProfile” tool. The cut forward and reverse reads (FastQ) were filtered, denoised and trimmed to 180 and 150 nucleotides, respectively, using the “filterAndTrim” tool. Overall, >92%, >75%, >97%, >71%, >96%, >91%, and >40% of input reads were retained and 0.03%, 0.01%, 0.58%, 0.01%, 0.01%, 0.00%, and 0.00% of merged paired reads were removed as chimeras for L42411, L21554, L45689, L9400, L20587, L15709_A, and L30575, respectively. For data cleaning purposes, the proportion (%) of each ASV in each sample was calculated. In an effort to remove misassigned reads caused by sample bleeding or barcode hopping, a threshold of <2.5% was applied (Mitra et al., 2015). This involved removing the reads of ASVs whose proportions represented <2.5% in a sample and replacing them with ‘0’. Alternative thresholds of <1% and <0.5% were tested but not selected for further downstream analysis. A reference table containing frequencies of sequence fragments of all 7 SNP loci was created, with options for each possible SNP polymorphism. This reference table was imported into DADA2 and the “AssignSpecies” tool was used to assign the remaining ASVs to a reference sequence if they were a perfect match. ASVs were imported into CLC Main Workbench (version 22.0) (CLC bio, Qiagen, Australia) and “BLAST at NCBI” was performed. Based on the BLAST results, ASVs that were not identified as *D. immitis* were excluded and ASV proportions for each sample were re-calculated. The remaining *D. immitis* ASVs were aligned to their respective SNP reference sequences and visually inspected using CLC Main Workbench.

### 2.11. Animal Ethics

Research performed at the University of Sydney was approved by the Animal Research Ethics Committee, which follows the NSW Animal Research Act 1985 and the Australian code for the care and use of animals for scientific purposes 8^th^ edition (2013).

### 2.12. Data accessibility

Raw FastQ sequences were deposited at SRA NCBI BioProject: PRJNA889241. Additional data for this study is available at LabArchives (https://dx.doi.org/10.25833/ygs4-ha97).

## 3. Results

### 3.1. Blood collection pre-and post-ML treatment

A total of 45 dogs (Dog 1 to Dog 45) and their 78 blood samples (HW01 to HW63, HW65 to HW79) were collected and followed up over 2.5 years (2019-2022) as suspect to be infected with *D. immitis* based on the veterinarians’ judgement (Supplementary Table S3). At the beginning of the study, infection with *D. immitis* was confirmed in 44/45 dogs using at least one diagnostic test. Dogs testing positive for *D. immitis* were domiciled at eleven locations across eastern Australia, with thirty-eight dogs from Queensland and six dogs from New South Wales (Figure 1). All dog bloods were processed using an exhaustive diagnostic laboratory workflow (Figure 2). Twenty-three (n=23) dogs were sampled at the time of diagnosis (pre-ML treatment) and at least once post-diagnosis; 14 dogs once after treatment (post-ML), eight dogs twice after treatment, and one dog (Dog 40) three times after treatment (Figure 2, 3). In addition, twenty-two dogs were only sampled pre-ML treatment and following samples were not possible to obtain.

**Fig. 1.**
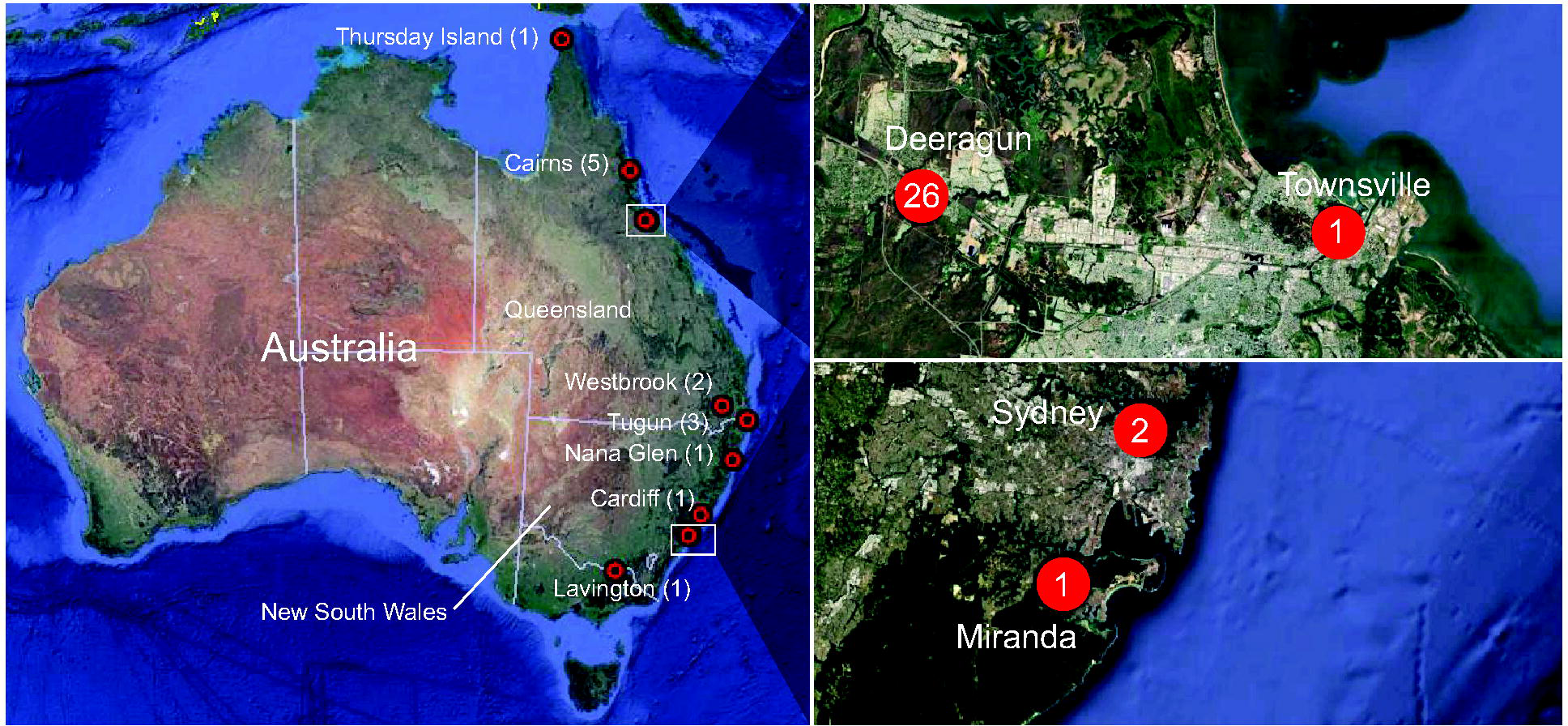
Location of dogs infected with *Dirofilaria immitis* included in this study from Eastern Australia. These dogs (n=44) were sampled between 2019-2022 for presence of *D. immitis* antigen, *D. immitis* DNA and associated *Wolbachia* DNA. The red circles indicate a location that is associated with the town name and the number indicates the number of *D. immitis*-positive dogs using any test. On the right, two close-ups of Townsville, Queensland and Greater Sydney, New South Wales are shown. Image from Google Earth; data from SIO, NOAA, U.S Navy, NGA, GEBCO (Image Landsat/Copernicus, CNES/Airbus, Maxar Technologies).

**Fig. 2.**
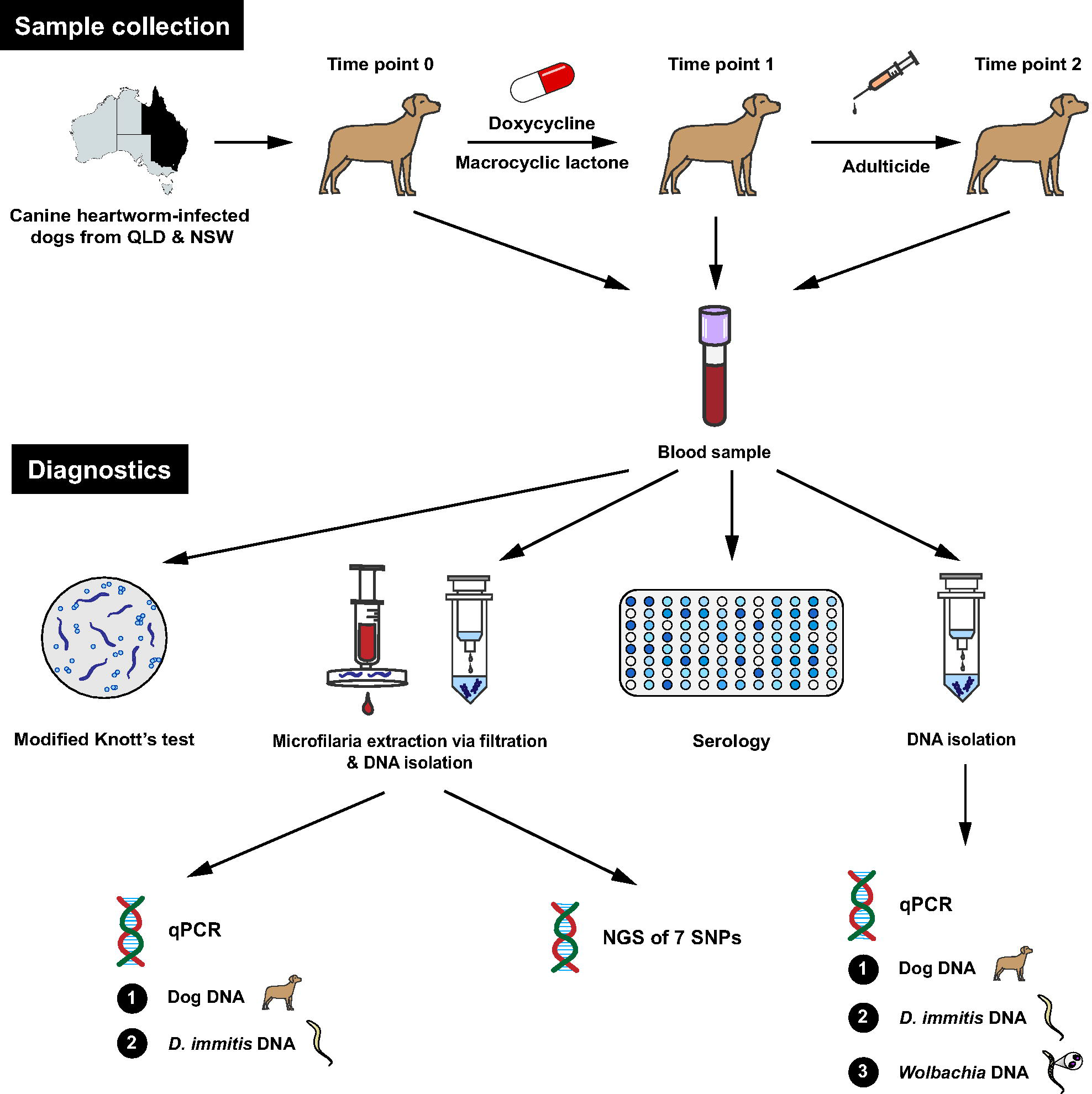
Diagnostic methods for the diagnosis of *Dirofilaria immitis* infection and response to macrocyclic lactone administration in dogs. Blood samples were collected at the time of diagnosis, followed by the administration of doxycycline antibiotic and macrocyclic lactone, and then the administration of adulticide injections. Each blood sample was assayed using a Modified Knott’s test to quantify microfilariae, microfilariae were extracted from blood via filtration for DNA isolation, plasma samples were assayed for the presence of *D. immitis* antigen and DNA isolated directly from blood was assayed quantitatively for the presence of *D. immitis* DNA and endosymbiotic *Wolbachia* DNA. DNA isolated from filtered microfilariae were tested for the presence of *D. immitis* DNA and endosymbiotic *Wolbachia* DNA in addition to single nucleotide polymorphisms (SNPs) associated with macrocyclic lactone resistance from the USA. SNPs were assayed using Illumina amplicon metabarcoding next generation sequencing (NGS).

### 3.2. Effects of ML administration on *Dirofilaria immitis* microfilariae counts

A total of 37/44 (84.1%) *D. immitis*-positive dogs were microfilaraemic at the beginning of the study, with a mean initial microfilariae count of 7,689 microfilariae/mL. Sequential Modified Knott’s tests were performed to demonstrate the suspect ML-resistance phenotypes of *D. immitis* (Figure 2). The percentage (%) reduction of microfilariae counts after ML administration was obtained for 22 dogs that had sequential blood samples. The mean initial microfilariae count for 21 dogs was 6,617 microfilariae/mL (min. 38 to max. 38,250 microfilariae/mL) plus one had 1 microfilaria/mL (Dog 1) (Figure 3, Table 1). In addition, one dog (Dog 19) had 0 microfilariae/mL, but tested positive on PCR and serology so a post-ML sample was also obtained and processed. Sixteen dogs (73%, 16/22) had ≤ 90% reduction in microfilariae count, including three dogs (Dog 25, Dog 2, Dog 7) which had an increase in microfilariae (Figure 3, Table 1).

**Fig. 3.**
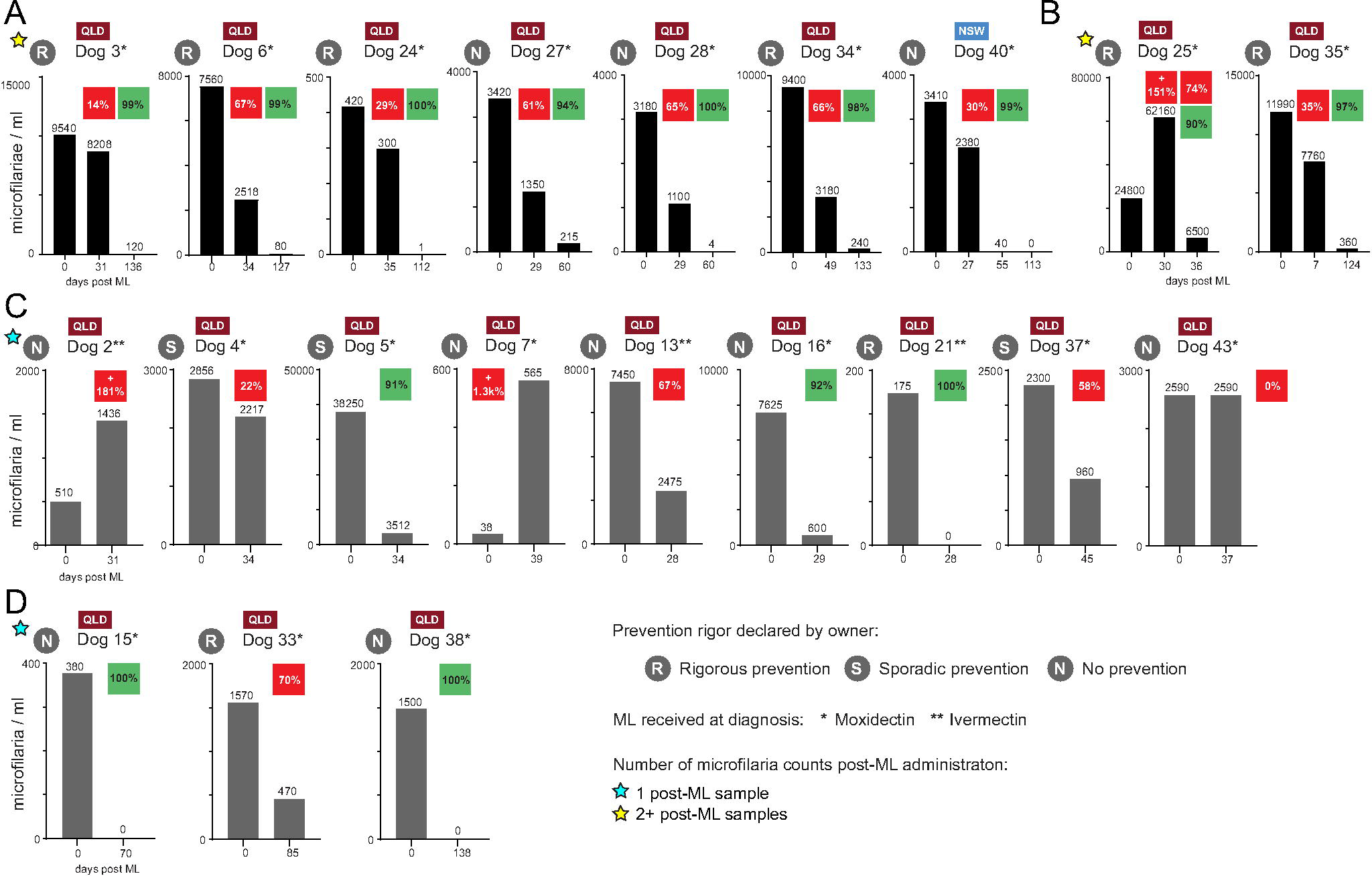
Sequential Modified Knott’s tests performed on on 21 dogs positive for *Dirofilaria immitis.* (A) Dogs (n=7) that exhibited <90% (red square %) microfilarial reduction at the first post-ML sampling, but consecutive second post-ML sampling demonstrated >90% microfilarial reduction (green square %). (B) Dog 25 at the first post-ML sampling had an increased number of microfilariae, but one week later the microfilariae were reduced by <90% (red squares); however, if comparing the 1^st^ and 2^nd^ post-ML samples, the reduction was >90%. Dog 35 has had its 1^st^ sample post-ML collected on day 7. (C) Dogs (n=9) that were sampled only once post-ML with variable reduction of microfilariae. (D) Dogs (n=3) that were sampled once, yet this sampling was delayed to 70 or more days post-ML. QLD – Queensland, Australia, NSW – New South Wales, Australia, yellow star – dogs with at least two consecutive post-ML blood samples tested, ML – macrocyclic lactone.

**Table 1.** Microfilariae quantification, antigen and qPCR diagnostics on Dirofilaria immitis infected dogs from Australia.

| Dog | Location<br>(Australia) | *D. immitis* prevention |  | Microfilariae reduction |  |  | DiroCHEK®<br>ELISA |  | qPCR C<sub>t</sub> |  |
| --- | --- | --- | --- | --- | --- | --- | --- | --- | --- | --- |
|  |  | Compliance<br>(past 12<br>months) | ML at<br>diagnosis | pre-ML/post-ML |  | % mf<br>change | pre-ML/post-ML |  | pre-ML/post-ML |  |
|  |  |  |  | ID | mf/ml |  | Neat | Heat | *D.*<br>*immitis* | *Wolbachia* |
| >90% microfilariae (mf) reduction |  |  |  |  |  |  |  |  |  |  |
| 1 | Cairns, QLD | Sporadic | HeartGard Plus | HW1/HW2 | 1/0 | -100 | +/- | +/+ | 37.7/- | -/- |
| 15 | Deeragun, QLD | No | ProHeart SR-12 | HW25/HW36 | 380/0 | -100 | +/- | +/+ | 26.0/- | -/- |
| 21 | Tugun, QLD | Yes | HeartGard Plus | HW31/HW34 | 175/0 | -100 | -/+ | +/+ | 32.2/32.1 | -/- |
| 38 | Deeragun, QLD | n/a | ProHeart SR-12 | HW63/HW75 | 1,500/0 | -100 | +/+ | +/+ | 23.4/- | 29.5/- |
| 16 | Deeragun, QLD | n/a | ProHeart SR-12 | HW26/HW33 | 7,625/600 | -92 | +/+ | +/+ | 23.1/25.3 | 28.1/34.7 |
| 5 | Deeragun, QLD | Sporadic | ProHeart SR-12 | HW7/HW13 | 38,250/3,512 | -91 | +/+ | +/+ | 20.0/22.4 | 25.1/32.3 |
| <90% microfilariae (mf) reduction |  |  |  |  |  |  |  |  |  |  |
| 33 | Deeragun, QLD | Yes | ProHeart SR-12 | HW55/HW59 | 1,570/470 | -70 | +/+ | +/+ | 24.6/25.2 | 30.3/- |
| 13 | Cairns, QLD | No | HeartGard Plus | HW20/HW22 | 7,450/2,475 | -67 | +/+ | +/+ | 20.5/21.8 | 25.2/32.8 |
| 6 | Deeragun, QLD | Yes | ProHeart SR-12 | HW8/HW15 | 7,560/2,518 | -67 | +/+ | +/+ | 21.3/22.3 | 26.7/32.0 |
| 34 | Deeragun, QLD | Yes | ProHeart SR-12 | HW56/HW58 | 9,400/3,180 | -66 | +/+ | +/+ | 21.2/22.9 | 27.3/35.5 |
| 28 | Westbrook, QLD | No | ProHeart SR-12 | HW44/HW49 | 3,180/1,100 | -65 | +/+ | +/+ | 25.1/24.9 | 28.8/33.7 |
| 27 | Westbrook, QLD | No | ProHeart SR-12 | HW43/HW48 | 3,420/1,350 | -61 | +/+ | +/+ | 25.8/23.9 | 29.8/33.8 |
| 37 | Deeragun, QLD | Sporadic | ProHeart SR-12 | HW61/HW65 | 2,300/960 | -58 | +/+ | +/+ | 22.6/25.4 | 29.9/- |
| 35 | Deeragun, QLD | Yes | ProHeart SR-12 | HW57/HW62 | 11,990/7,760 | -35 | +/+ | +/+ | 19.5/22.7 | 27.3/32.6 |
| 40 | Nana Glen, NSW | No | ProHeart SR-12 | HW68/HW71 | 3,410/2,380 | -30 | +/- | +/+ | 22.7/27.0 | 28.2/- |
| 24 | Deeragun, QLD | Yes | ProHeart SR-12 | HW38/HW40 | 420/300 | -29 | +/+ | +/+ | 26.6/26.0 | 31.7/33.5 |
| 4 | Deeragun, QLD | Sporadic | ProHeart SR-12 | HW6/HW12 | 2,856/2,217 | -22 | +/+ | +/+ | 22.0/22.9 | 26.6/32.3 |
| 3 | Deeragun, QLD | Yes | ProHeart SR-12 | HW5/HW14 | 9,540/8,208 | -14 | +/+ | +/+ | 22.7/24.4 | 26.1/35.1 |
| 43 | Deeragun, QLD | n/a | ProHeart SR-12 | HW74/HW78 | 2,590/2,590 | 0 | +/+ | +/+ | 24.7/22.7 | 30.6/33.1 |
| 19 | Tugun, QLD | Yes | HeartGard Plus | HW29/HW35 | 0/0 | n/a | +/- | +/+ | 33.1/- | -/- |
| Increase in microfilariae (mf) |  |  |  |  |  |  |  |  |  |  |
| 7 | Townsville, QLD | n/a | ProHeart SR-12 | HW9/HW16 | 38/565 | +1,386 | +/+ | +/+ | 28.7/24.0 | 31.8/34.3 |
| 2 | Thursday Is, QLD | No | HeartGard Plus | HW3/HW4 | 510/1436 | +182 | +/+ | +/+ | 21.8/22.9 | 27.0/31.9 |
| 25 | Deeragun, QLD | Yes | ProHeart SR-12 | HW39/HW41 | 24,800/6,2160 | +151 | +/+ | +/+ | 21.3/24.1 | 26.1/28.8 |

Note: n/a – not available, + (DiroCHEK) – positive, - (DiroCHEK) – negative, - (qPCR) – negative (>40), ML – macrocyclic lactone, mf – microfilariae, HeartGard Plus (ivermectin/pyrantel; Boehringer-Ingelheim), ProHeart SR-12 (moxidectin; Zoetis).

Using seven dogs sampled pre-ML which were then administered moxidectin (ProHeart SR-12, s.c.) and sampled again at least twice post-ML, we show a 47% (14%-67%, Figure 3A) reduction of microfilariae by day 27-49. This was reduced to 95% (73%-100%)by 60-136 days. Four of these dogs were subjectively considered to be on ‘rigorous’ heartworm prevention by their owners (Figure 3A). All remaining dogs but three were administered moxidectin (ProHeart SR-12, s.c.) at the time of diagnosis (Figure 3B,C,D). Three dogs were administered ivermectin (HeartGard Plus, p.o.). Dog 21 on ivermectin (HeartGard Plus) completely reduced their microfilariae by day 28 from a relatively low initial microfilaremia of 175 microfilariae/mL (Figure 3C). The remaining two dogs (Dog 2, 13) on ivermectin (HeartGard Plus) experienced <90% reductions in microfilaremia post-ML, with Dog 2 experiencing an increase in microfilariae by 181% (Figure 3C).

### 3.3. Presence of heartworm antigen using DiroCHEK**^®^** ELISA

Out of all dogs with confirmed *D. immitis* infection at the start of this study, 41/44 (93.2%) dogs initially tested positive for *D. immitis* antigen using DiroCHEK^®^ ELISA with unheated (neat) plasma samples and 43/44 (97.7%) initially tested positive using DiroCHEK^®^ ELISA with heated plasma samples. Out of all blood samples collected in this study, 88.2% (67/76) of neat plasma samples and 98.7% (75/76) of heated plasma samples tested positive for *D. immitis* antigen using DiroCHEK^®^ ELISA, respectively (Figure 4A, Table 1). A paired t-test was performed on the raw DiroCHEK^®^ ELISA data, revealing a significant difference in OD_620_ between neat and heated plasma samples (Figure 4B). Moderate positive correlations were found between raw microfilariae counts and the OD_620_ of neat and heated dog plasma in DiroCHEK^®^ ELISA (Figure 5A). When performing the modified Knott’s test on samples with microfilariae counts >20,000 mf/mL, we observed the overlap and clumping of microfilariae, which made counting these samples difficult. To determine whether similar statistical results were still obtained for samples with microfilariae counts <20,000 mf/mL, the correlation analyses between microfilariae counts and the DiroCHEK^®^ ELISA were repeated. By doing so, moderate positive correlations were maintained (Figure 5B).

**Fig. 4.**
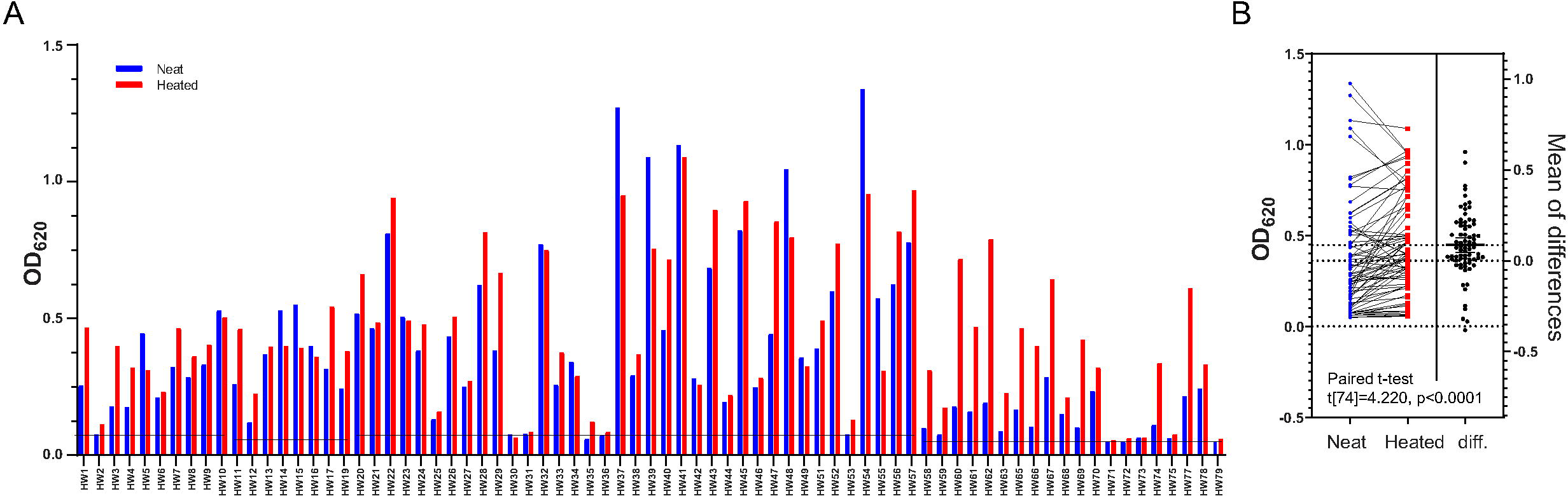
*Dirofilaria immitis* antigen testing (DiroCHEK^®^ ELISA, Zoetis, Australia) on 76 blood samples collected from 44 dogs in Australia. (A) Neat plasma samples from 41/44 dogs and heated plasma samples from 43/44 dogs initially tested positive for *D. immitis* antigen. Out of all blood samples tested, 67/76 neat samples and 75/76 heated samples were positive for *D. immitis* antigen. A threshold of 3×SD (Mean[optical density of negative controls]) was applied to each run to distinguish between *D. immitis*-negative and *D. immitis*-positive samples. Thresholds of 0.0784 (HW1-HW10), 0.0596 (HW11-HW17, HW19), 0.0786 (HW20-HW49, HW51-HW57), and 0.0532 (HW58-HW63, HW65-HW75, HW77-HW79) were applied. (B) Paired t-test on DiroCHEK^®^ ELISA data demonstrated a significant difference in optical density between neat and heated plasma samples (Paired t-test, t[74]= 4.220, p <0.0001). Data analysis and graphs were produced using GraphPad Prism (version 9.3.1).

**Fig. 5.**
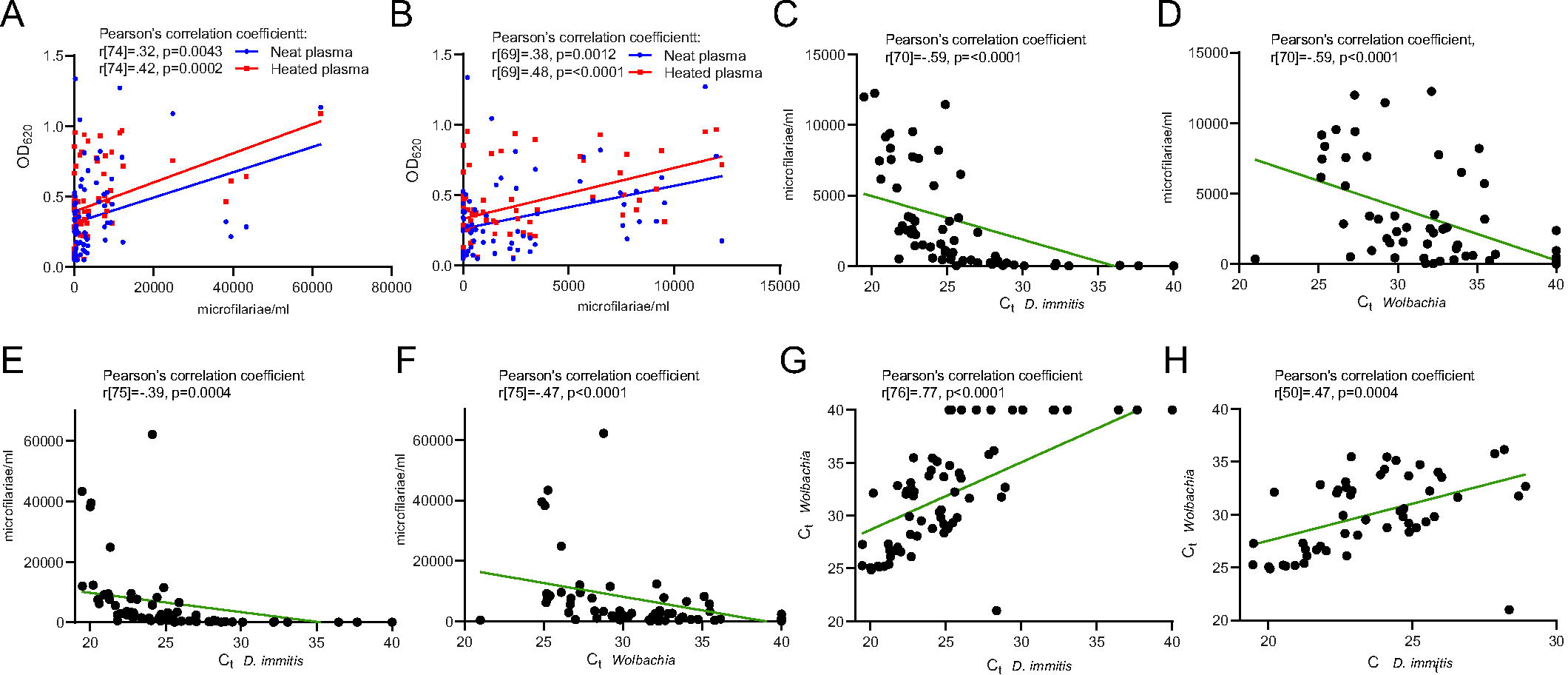
Correlations between microfilariae count (mf/mL), DiroCHEK^®^ ELISA (Zoetis, Australia) of neat and heated plasma, and qPCR of whole blood DNA targeting *D. immitis* and associated *Wolbachia*. (A) Moderate positive correlations between microfilariae counts and OD_620_ of neat (n=76) and heated (n=76) plasma. (B) Moderate positive correlations between microfilariae counts and OD_620_ of neat (n=71) and heated (n=71) plasma for samples with microfilariae counts <20,000 mf/mL. (C) Strong negative correlation between *D. immitis* qPCR C_t_-values of whole blood DNA and microfilariae counts (n=72) for samples with <20,000 mf/mL. (D) Strong negative correlation between *Wolbachia* qPCR C_t_-values of whole blood DNA and microfilariae counts (n=72) for samples with <20,000 mf/mL. (E) Moderate negative correlation between *D. immitis* qPCR C_t_-values of whole blood DNA and microfilariae counts (n=77). (F) Moderate negative correlation between *Wolbachia* qPCR of whole blood DNA and microfilariae counts (n=77). (G) Strong positive correlation between *D. immitis* and *Wolbachia* qPCR C_t_-values of whole blood DNA (n=78). (H) Moderate positive correlation between *D. immitis* and *Wolbachia* qPCR C_t_-values of whole blood DNA (n=52) for samples with C_t_-values <40. Data analysis and graphs were produced using GraphPad Prism (version 9.3.1).

### 3.4. Presence of Dirofilaria immitis and Wolbachia sp. of D. immitis using qPCR

DNA samples obtained from dog blood were subjected to qPCR targeting the *cox1* gene of *D. immitis* and the *ftsZ* gene of *D. immitis*-associated *Wolbachia*. *D. immitis* and *Wolbachia* DNA were initially detected in 39/44 (88.6%) and 34/44 (77.3%) dogs with confirmed *D. immitis* infection, respectively. At Day 0 (pre-ML), *D. immitis* and *Wolbachia* were detected in 100% (23/23, C_t_ mean: 24.63, C_t_ range: 19.48-37.68) and 82.61% (19/23, C_t_ mean: 28.21, C_t_ range: 25.07-31.75) of dogs, respectively. At ∼Day 30 (post-ML), *D. immitis* was detected in 82.61% (19/23, C_t_ mean: 24.37, C_t_ range: 21.80-32.13) of dogs and *Wolbachia* was detected in 65.22% (15/23, C_t_ mean: 33.10, C_t_ range: 28.78-35.48) of dogs. By Day 30+, 77.8% (7/9, C_t_ mean: 29.00, C_t_ range: 25.89-36.45) of dogs were positive for *D. immitis* and 33.33% (3/9, C_t_ mean: 30.27, C_t_ range: 21.01-35.78) of dogs were positive for *Wolbachia*. The *D. immitis*-specific qPCR on extracted microfilariae DNA found one additional *D. immitis* DNA positive dog (Dog 31: 0 microfilariae/mL) that tested negative for *D. immitis* DNA on a sample from whole blood. In addition, two dogs (Dog 1: 1 microfilaria/mL, Dog 20: 0 microfilariae/mL) were *D. immitis* DNA positive only on whole blood but *D. immitis* DNA-negative using extracted microfilariae (Supplementary Table S3).

A strong negative correlation was demonstrated between the *D. immitis* qPCR C_t_-values of whole blood DNA and microfilariae counts that were <20,000 mf/mL (Figure 5C). Samples with microfilariae counts >20,000 mf/mL were excluded from this analysis due to difficulties with counting clumped and overlapping microfilariae. Similarly, a strong negative correlation was observed between the *Wolbachia* qPCR C_t_-values of whole blood DNA and microfilariae counts <20,000 mf/mL (Figure 5D). These statistical analyses were repeated to determine whether similar correlations were still obtained when including samples with microfilariae counts > 20,000 mf/mL. By doing so, a moderate negative correlation was shown between the *D. immitis* qPCR C_t_-values of whole blood DNA and microfilariae counts (Figure 5E). For all *Wolbachia* qPCR of whole blood DNA, a moderate negative correlation was observed with microfilariae counts (Figure 5F). A strong positive correlation was found between the *D. immitis* and *Wolbachia* qPCR C_t_-values of whole blood DNA samples (Figure 5G). Samples with C_t_-values of 40 were considered negative and excluded from the analysis, resulting in a moderate positive correlation (Figure 5H).

Paired t-tests of the raw *D. immitis* and *Wolbachia* qPCR data revealed statistically significant increases in C_t_-value from pre-ML to first post-ML sampling (Figure 6A,B). To determine whether similar results were obtained for samples with pre-ML C_t_-values <30 in the *D. immitis* qPCR, the analyses were repeated. By doing so, we confirmed a significant increase in C_t_-value. (*D. immitis* paired t-test, t[19]=2.159, p=0.0439). To determine if we would obtain similar results for samples with pre-ML C_t_-values <35 in the *Wolbachia* qPCR, the analyses were repeated. This confirmed a significant increase in C_t_-value. (*Wolbachia* qPCR paired t-test, t[18]=9.290, p <0.0001). Similarly, paired t-tests of the raw *D. immitis* and *Wolbachia* qPCR data showed statistically significant increases in C_t_-values from pre-ML to second post-ML sampling (Figure 6C,D).

**Fig. 6.**
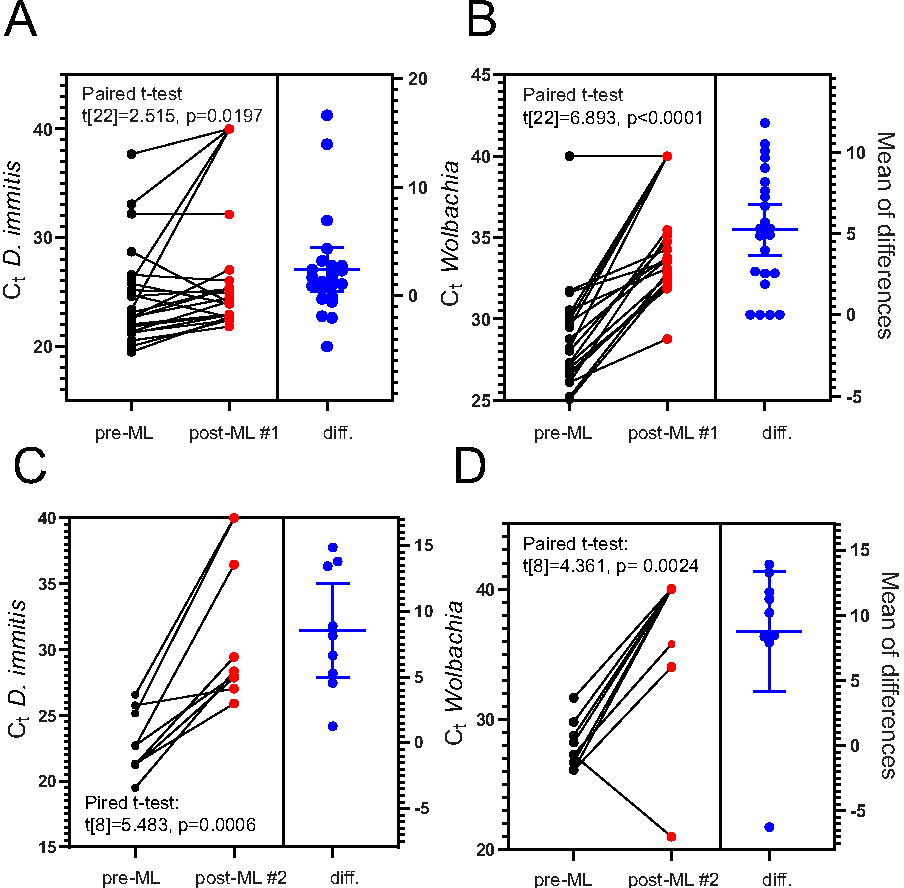
Paired t-test analyses on *D. immitis* and *Wolbachia* qPCR C_t_-values pre- and post-ML treatment. (A) *D. immitis* qPCR demonstrated statistically significant increase in C_t_-value between pre-ML and first post-ML sampling (n=23). (B) *Wolbachia* qPCR demonstrated statistically significant increase in C_t_-value between pre-ML and first post-ML sampling (n=23). (C) *D. immitis* qPCR demonstrated statistically significant increase in C_t_-value between pre-ML and second post-ML sampling (n=9). (D) *Wolbachia* qPCR demonstrated statistically significant increase in C_t_-value between pre-ML and second post-ML sampling (n=9). Data analysis and graphs were produced using GraphPad Prism (version 9.3.1).

### 3.5. Illumina PCR amplicon NGS identification of ML-susceptible *D. immitis* in Australian dogs

Illumina NGS amplicon metabarcoding from 64 extracted microfilariae DNA samples from 37 dogs and one DNA sample from an adult *D. immitis* (P6/20D, positive control) was performed to determine the nucleotide proportions at 7 SNPs of interest (65 DNA samples×7 loci = 455 PCRs) (Figure 7). Successful amplification of the target region was achieved for 96.5% (439/455) of PCR samples. Three DNA samples had loci which failed the first-stage PCR amplification and were not submitted for Illumina NGS amplicon sequencing (HW34: 1/7 SNPs submitted, 0 microfilariae/mL; HW50: 5/7 SNPs submitted, 1 microfilariae /mL; HW51: 1/7 SNPs submitted, 0 microfilariae/mL). Only 5/7 SNPs were submitted for P6/20D. FastQ files were obtained, and SNP calling was successful for 98.2% (431/439) of submitted PCR samples. Four DNA samples had loci which failed to produce reads for SNP calling (HW29: 6/7 SNPs successful; HW34: 0/1 SNPs successful; HW50: 0/5 SNPs successful; HW51: 0/1 SNPs successful). For the purpose of reproducibility, an additional 28 PCR samples were submitted as ‘repeats’. These repeat samples demonstrated identical outcomes in terms of SNP calling. Overall, a total of 8,742,348 merged non-chimeric paired reads were obtained to acquire amplicon sequence variants (ASVs) which were matched to the reference ML ‘susceptible’ and ML ‘resistant’ allele for their respective SNP (Figure 7).

**Fig. 7.**
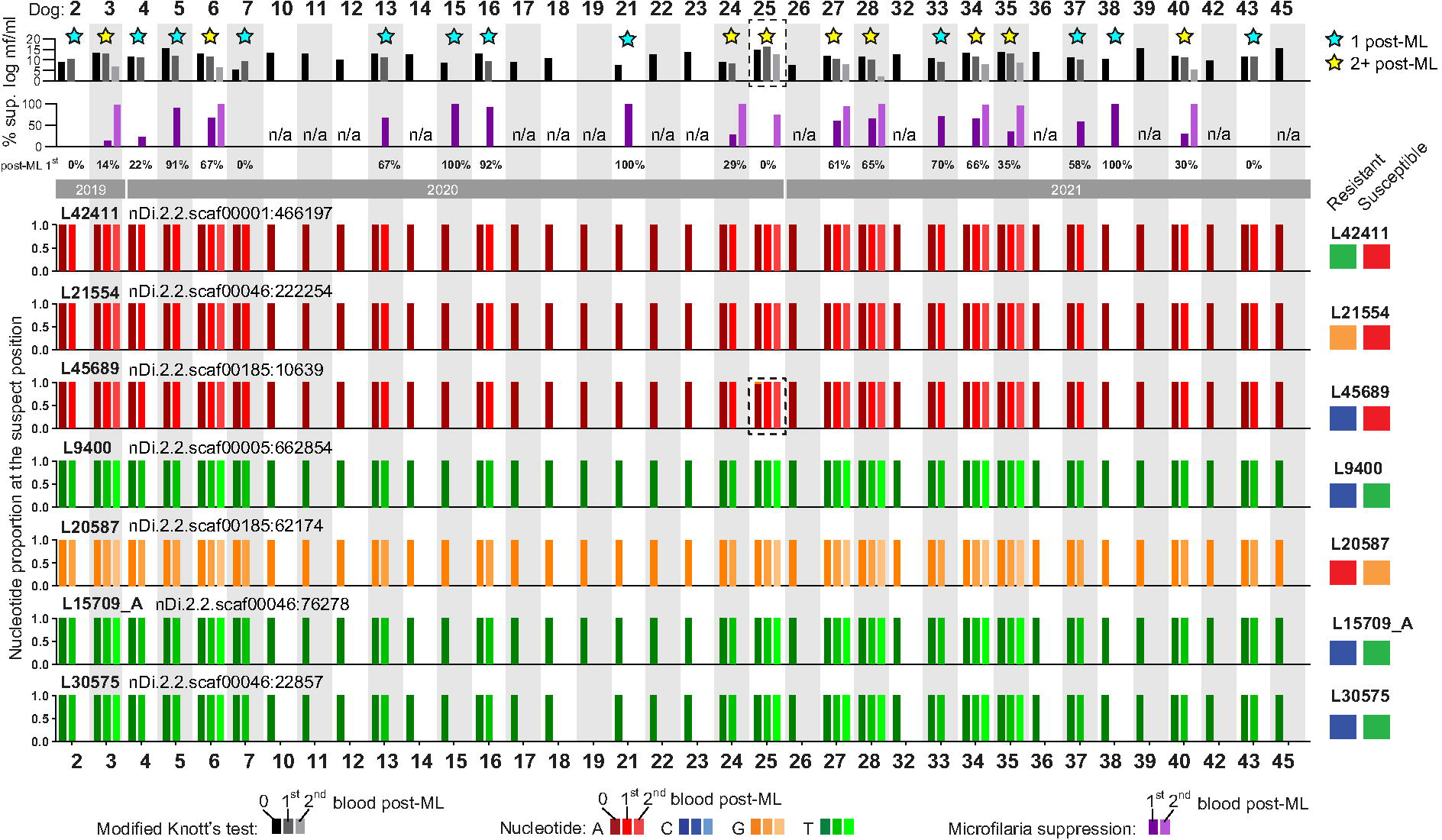
Australian dogs infected with *Dirofilaria immitis* and response to ML treatment. Dog identification number is indicated at the top. Upper-most bar chart represents log transformed microfilariae (mf) counts pre-ML and post-ML (black to grey; Modified Knott’s test). Yellow stars – dogs with at least two consecutive post-ML blood samples tested (2+ post-ML), blue stars – dogs with one post-ML blood sample tested (1 post-ML). Percent (%) reduction of microfilariae at the first and second post-ML time point is plotted (purple bars, n/a – not applicable due to absence of sequential sample). Percentage (%) reduction at 1^st^ post-ML timepoint is written along the year these samples were collected. Nucleotide proportions at SNP regions (identified according to nDi2.2. genome assembly contigs) are shown for all samples analysed. The blue, red, and orange represent the nucleotides, with the reference nucleotides associated with ML-resistance indicated for each SNP on the right. Dog 25 has microfilariae count and L45689 highlighted by dotted line rectangles.

#### 3.5.1. L42411 (Di.2.2.scaf00001:466197)

A total of 1,368,512 merged non-chimeric paired reads were obtained for L42411 with an average of 20,426 paired reads per sample. Overall, 62/65 DNA samples displayed 100% of the susceptible nucleotide at the suspect position. All 62 samples had 100% of their reads assigned to a single ASV (L42411_ASV1) which was a perfect match to the ML ‘susceptible’ reference sequence.

#### 3.5.2. L21554 (nDi.2.2.scaf00046:222254)

A total of 1,167,570 merged, non-chimeric paired reads for 362 ASVs were obtained for L21554 with an average of 17,690 paired reads per sample. Overall, 62/65 DNA samples displayed 100% of the susceptible nucleotide at the suspect position. Of these 62 samples, 14 had 100% of their reads assigned to a single ASV (L21554_ASV1) which was a perfect match to the ‘susceptible’ reference sequence. The remaining 48 samples contained L21554_ASV1 along with low levels of other ASVs (L21554_ASV2, L21554_ASV3) which possessed a SNP at another position in the sequence.

#### 3.5.3. L45689 (nDi.2.2.scaf00185:10639)

A total of 1,664,418 merged, non-chimeric paired reads for 101 ASVs were obtained for L45689 with an average of 23,777 paired reads per sample. Overall, 61/65 DNA samples had 100% of the susceptible nucleotide at the suspect position. Of these 61 samples, 15 had 100% of their reads assigned to a single ASV (L45689_ASV1) which was a perfect match to the ‘susceptible’ reference sequence. The remaining 46 samples contained multiple ASVs (L45689_ASV1, L45689_ASV2, L45689_ASV3, L45689_ASV4, L45689_ASV8, L45689_ASV10, L45689_ASV11) which possessed 1-2 SNPs at other positions in the sequence. One sample (Dog 25 – HW39) contained low levels (3.8%) of L45689_ASV6 which possessed a nucleotide of unknown ML association at the suspect position (Figure 8).

**Fig 8.**
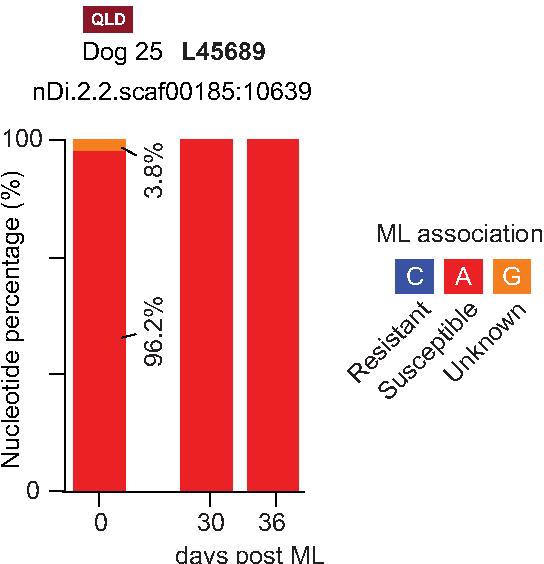
Nucleotide percentages at a SNP region (L45689) previously associated with ML-resistance in the USA for Dog 25. Blood samples were collected at 0-, 30- and 36-days post-ML treatment and DNA was isolated from filtered *Dirofilaria immitis* microfilariae. The blue, red, and orange boxes in the key represent the nucleotides and their associated ML-resistance status.

#### 3.5.4. L9400 (nDi.2.2.scaf00005:662854)

A total of 908,148 merged, non-chimeric paired reads for 543 ASVs were obtained for L9400 with an average of 12,613 paired reads per sample. Overall, 62/65 DNA samples displayed 100% of the susceptible nucleotide at the suspect position. Of these 62 samples, 57 had 100% of their reads assigned to a single ASV (L9400_ASV1) which was a perfect match to the ‘susceptible’ reference sequence. The remaining 5 samples contained L9400_ASV1 along with other ASVs (L9400_ASV2, L9400_ASV30) which possessed a SNP at another position in the sequence.

#### 3.5.5. L20587 (nDi.2.2.scaf00185:62174)

A total of 1,600,533 merged, non-chimeric paired reads for 90 ASVs were obtained for L20587 with an average of 23,537 paired reads per sample. Overall, 62/65 DNA samples displayed 100% of the susceptible nucleotide at the suspect position. Of these 62 samples, 57 had 100% of their reads assigned to a single ASV (L20587_ASV1) which was a perfect match to the ‘susceptible’ reference sequence. The remaining five samples contained L20587_ASV1 along with other ASVs (L20587_ASV2, L20587_ASV4, L20587_ASV5, L20587_ASV6, L20587_ASV8, L20587_ASV11) which possessed a SNP at another position in the sequence.

#### 3.5.6. L15709_A (nDi.2.2.scaf00046:76278)

A total of 1,526,468 merged, non-chimeric paired reads for 78 ASVs were obtained for L15709_A with an average of 24,620 paired reads per sample. Overall, 61/65 DNA samples displayed 100% of the susceptible nucleotide at the suspect position. All 61 samples had 100% of their reads assigned to a single ASV. Thirty samples had all reads assigned to L15709A_ASV1 which was a perfect match to the ‘susceptible’ reference sequence. The other 31 samples had all reads assigned to L15709A_ASV2 which possessed a deletion elsewhere in the sequence.

#### 3.5.7. L30575 (nDi.2.2.scaf00046:22857)

A total of 506,699 merged, non-chimeric paired reads for 542 ASVs were obtained for L30575 with an average of 8,173 paired reads per sample. Overall, 60/65 DNA samples displayed 100% of the susceptible nucleotide at the suspect position. All 60 samples had 100% of their reads assigned to a single ASV (L30575_ASV1) which was a perfect match to the ‘susceptible’ reference sequence.

## 4. Discussion

The use of MLs over the past 30 years has successfully reduced the prevalence of canine heartworm infection (Orr et al., 2020; Lau et al., 2021). However, the re-emergence of *D. immitis* infection in Queensland, Australia and the confirmation of ML-resistant isolates in the USA poses an important question of whether ML-resistance exists in Australia (Blagburn et al., 2011; Pulaski et al., 2014; Bourguinat et al., 2015; Blagburn et al., 2016; Nguyen et al., 2016; Maclean et al., 2017; McTier et al., 2017b; Orr et al., 2020; Panetta et al., 2021). By considering microfilariae counts as the phenotype and genotypic tests adopted from the USA, this study did not unequivocally demonstrate the presence of ML-resistant *D. immitis* in Australia. Although the sequential Modified Knott’s tests showed delayed clearance of circulating microfilariae compared to the > 90% reduction at 28 days described by McTier et al. (2017a), suggestive of suspect ML-resistance, no genotypic evidence was discovered. Our approach was in part modelled to that of Ballesteros et al. (2018), where 27 dogs underwent ‘microfilarial suppression testing’ *sensu* Geary et al. (2011) and SNP analysis with moxidectin (Elanco, Advantage Multi for dogs, spot on). By doing so, they validated the association between ML microfilaricidal response phenotype and SNP loci genotype (Ballesteros et al., 2018). The current study obtained a comparable sample size to Ballesteros et al. (2018), and adopted their 5-SNP predictive model which had 94.1% sensitivity and 100% specificity. We utilised Illumina amplicon metabarcoding coupled with DADA2 analysis to detect the susceptible and resistant SNPs, which provided us with reproducible results directly comparable to Ballesteros et al. (2018).

To treat canine heartworm disease, infected dogs are administered doxycycline and ML at the time of diagnosis (Day 0) to target the *Wolbachia* endosymbiont of *D. immitis*, prevent further infection by eliminating *D. immitis* L3/L4 and if possible (depending on the specific ML used) decrease circulating microfilariae (Nelson et al., 2020). To target the adult stages of *D. immitis*, clinicians in Australia typically follow the three-dose protocol recommended by the American Heartworm Society (Nelson et al., 2020). Importantly, for over 10 years, Australian veterinarians had access to a yearly injectable moxidectin (ProHeart SR-12, Zoetis) preventative (Mwacalimba et al., 2021). ProHeart SR-12 does not have a claim for circulating microfilariae in Australia, but it has been shown to reduce circulating microfilariae by >90% 28 days post-administration (McTier et al., 2017a).

The utilised sequential Modified Knott’s test to calculate the reduction is not identical to the ‘microfilarial suppression testing’ *sensu* Geary et al. (2011) with spot-on moxidectin in the USA based on the label claim for microfilariae and 99% clearance within 14 days in an experimental trial (Geary et al., 2011; Bowman et al., 2015). We extended the post-ML interval to ∼30 days for two reasons. Firstly, to suit the veterinary practice when owners brought their dog back for a health check-up, and secondly to utilise the published microfilaricidal effect of ProHeart SR-12 (McTier et al., 2017a). While the choice of ML-preventative was at the veterinarian’s discretion, most were injected with ProHeart SR-12 in our study. The failure of 16 dogs (ML at diagnosis; moxidectin microspheres: 14 dogs, ivermectin: 2 dogs) to reduce microfilaremia by >90% was considered phenotypic evidence of suspect ML-resistance. This finding was not corroborated by genetic SNP analysis or phenotypic testing beyond 30 days, as dogs that were re-tested eventually reached a >90% reduction in microfilaremia by Day 55 to Day 136. Delays in reaching >90% microfilarial reduction following ML treatment have been previously observed. In McTier et al. (2017a), single doses of ProHeart 6 (Zoetis, moxidectin microspheres, 170 µg/kg) and oral moxidectin (250 µg/kg) on Day 0 resulted in ≤90% reductions in mean microfilariae counts for the ML-resistant isolate ZoeMO-2012 at Day 28 (ProHeart 6: 86.5%, oral moxidectin: 83.6%), whereas ProHeart 12 (Zoetis, moxidectin microspheres, equivalent to ProHeart SR-12 registered in Australia, 500 µg/kg) led to a 91.2% reduction in microfilariae. At Day 84, all three MLs reached >90% reductions in microfilariae (ProHeart 6: 96.8%, ProHeart 12: 96.9%, oral moxidectin: 92.6%) (McTier et al., 2017a). In contrast to the experimental trial by McTier et al. (2017a), the current study sampled heterogeneous privately-owned dogs which most likely carried a diverse population of *D. immitis*..

The variability of reduction in microfilariae counts post-ML administration in the present study can be attributed to the varying effectiveness of MLs to clear circulating microfilariae (Prichard, 2021). In Australia, no product has a label claim for microfilariae, unlike in the USA where Advantage Multi for dogs (10% imidacloprid + 2.5% moxidectin topical solution) has such a claim and was therefore used by Ballesteros et al. (2018) in their study (Bowman et al., 2015). A microfilaricidal effect was noted early on for ivermectin in dogs at a dose of 50 µg/kg, but due to toxicity in dolichocephalic breeds, this high dose is not on any label (Blair and Campbell, 1979). The current preventative dose against <30-day old *D. immitis* with ivermectin is only 6 µg/kg, which is considered insufficient to clear microfilariae efficiently. Milbemycin oxime at 500 µg/kg is considered microfilaricidal, but no registered product currently has such a claim (Prichard, 2021). Moxidectin in the spot-on formulation (Elanco, Advantage Multi for dogs/Advocate for dogs at a minimum dose of 2,500 µg/kg) was demonstrated to be effective against microfilariae, which has been claimed as such in the Advantage Multi for dogs (Elanco) marketed in the USA (McCall et al., 2014; Bowman et al., 2015).

Sequential quantitative Modified Knott’s tests to enumerate the number of circulating *D. immitis* microfilariae pre-and post-ML treatment enables clinicians to determine whether moxidectin treatment is successfully clearing microfilaremia in the infected dog and whether the dog remains a threat to other dogs. However, performing a non-quantitative Knott’s test is considered impractical for clinicians as it can be time-consuming, labour-intensive, and prone to human error. Alternative tools that are more rapid and accurate are required to measure changes in post-treatment microfilaremia levels. The blood samples in this study were screened for *D. immitis* using DiroCHEK^®^ and qPCR targeting *D. immitis* and its associated *Wolbachia* endosymbiont. Based on our data, performing DiroCHEK^®^ on neat or heated plasma was not suitable for the assessment of ML-treatment efficacy due to its lower correlation with the Modified Knott’s test. The *D. immitis* and *Wolbachia* qPCRs produced higher correlations with the Modified Knott’s test and were able to demonstrate the success of ML-treatment for reducing microfilariae levels. We therefore suggest clinicians to use these faster molecular tools as a substitute for the Modified Knott’s test when monitoring the success of MLs in canine heartworm cases. The *D. immitis*-specific qPCR detected a larger proportion of infected dogs in this study compared to the *Wolbachia*-specific qPCR and may therefore be a better tool for measuring microfilariae levels. Future studies will benefit from a triplex qPCR targeting *D. immitis*, its associated *Wolbachia*, and dog DNA to reduce time and cost. The same *Wolbachia* qPCR assay was recently used to monitor the *Wolbachia* levels in dogs from Italy following doxycycline and Advocate (moxidectin) treatment (Laidoudi et al., 2020; Louzada-Flores et al., 2022). By doing so, they found that all dogs tested negative for *Wolbachia* after one month. Contrastingly, the current study saw only 8/28 dogs test negative for *Wolbachia* approximately 30 days after ML-treatment, with 4 of these dogs already being negative on Day 0. When infected dogs were re-sampled on Day 30+, only 6/9 tested negative for *Wolbachia*.

The SNPs analysed in Ballesteros et al. (2018) were able to distinguish between ML-susceptible and ML-resistant *D. immitis* in the USA. However, in the current study and Lau et al. (2021), no evidence of ML-resistance in Australian *D. immitis* was found using these same ML resistance-associated SNPs. In Lau et al. (2021), the genome data of five adult *D. immitis* exhibited marked genetic differences compared to those previous phenotypically characterised *D. immitis* isolates from the USA. Genomics data from ML-susceptible and ML-resistant *D. immitis* are required to associate and specifically predict the ML-resistance in clinical cases in Australia. This will enable the early detection of ML resistance in Australia, allowing clinicians, researchers and the animal pharmaceutical industry to develop effective control measures to limit and manage the spread of resistant parasites.

Successful prevention of *D. immitis* heartworm can only be achieved with rigor in prevention due to inability to successfully remove older than 30-day *D. immitis* with current ML-based preventatives (Prichard, 2021). For veterinarians, however, the inquiry into rigor with ML-prevention will always be difficult with products administered by owners as they can unintentionally skip dosing if dogs regurgitate the dose (Atkins et al., 2014). An unintentionally missed ML-dose may lead to infection – apparent lack of efficacy (LOE) reports - despite owners’ assumed rigor with prevention. In our study, despite owners’ claims that dogs received ‘rigorous’ heartworm prevention with either moxidectin, milbemycin oxime or ivermectin for 12 months prior to the veterinarian’s diagnosis, 16 dogs tested positive for canine heartworm infection. Similar findings were encountered in Nguyen et al. (2016) where microfilaraemic dogs in Queensland previously received ‘rigorous’ prevention. In this study, owner compliance could not be confirmed and was solely based on the owner’s reports.

## 5. Conclusion

To confirm the presence of drug resistance in a parasite population, one must demonstrate that resistance is genetically inherited (Kohler, 2001). Since 2005, lack of efficacy complaints for heartworm preventatives led to speculation and controversy on the potential emergence of ML-resistant *D. immitis* in the USA (Hampshire, 2005; Atkins et al., 2014). It was not until 2014 that the first ML-resistant strain (LSU 10) was isolated in the USA and resistance to MLs was unequivocally confirmed (Pulaski et al., 2014). To do this, a laboratory dog was experimentally infected with L3s obtained from a field case, and then treated with ivermectin (Pulaski et al., 2014). Infective larvae from this dog were then successfully passaged into a second laboratory dog, which confirmed the genetic heritability of drug resistance (Pulaski et al., 2014). In Australia, we currently lack these types of specialised facilities where laboratory dogs can be experimentally infected with canine heartworm. Due to this limitation, it remains unknown whether ML-resistance is present in Australia, much like the 2005-2014 time period in the USA. Until we can demonstrate the heritability of ML-resistance in Australian *D. immitis* isolates via experimental infection studies, we can only speculate and test for resistance using the available tools as a proxy.

## Acknowledgements

We thank all the veterinary practitioners and owners of the patients, in particular Dr. Tony Phillis and his team at Greencross Deeragun for valiant effort to recruit patients and collect samples for screening. The work was in part supported by the Canine Research Foundation and Dogs Victoria, Australia and by the Australian Companion Animal Health Foundation Research Grant. RP is supported by an Australian Government Research Training Program Stipend and The Jean Walker Trust Fellowship. We thank Ilze Nel and Richard L’Estrange (Zoetis Australia Pty Ltd) for discussion around heartworm prevention.

## Supplementary Tables

**Supplementary Table S1:** Primers used throughout the study

**Supplementary Table S2:** SNPs in this study adopted from previous 5-SNPs models from the USA

**Supplementary Table S3:** Summary of Australian dogs infected with *Dirofilaria immitis* monitored over 2019-2022

