## Supplementary Table S1 for "Exploration of the sensitivity to macrocyclic lactones in the canine heartworm (*Dirofilaria immitis*) in Australia using phenotypic and genotypic approaches": MS_RoseHW_Supplementary_Table S1.docx

**Supplementary Table S1:** Primers used throughout the study

| **Assay specificity** | **Application** | **Target** | **Primer/probe** | **ID** | **Sequence (5'-3')** | **Reference** |
| --- | --- | --- | --- | --- | --- | --- |
| **Dog** | qPCR | ***GAPDH*** | Primer | S0631_DOG_F, S1072_DOG_F | TCAACGGATTTGGCCGTATTGG | (Orr et al., 2020; Panetta et al., 2021) |
|  |  |  |  | S0634_DOG_R, S1073_DOG_R | TGAAGGGGTCATTGATGGCG |  |
|  |  |  | Probe | S0632_DOG_P, S1074_DOG_P | CAGGGCTGCTTTTAACTCTGGCAAAGTGGA |  |
| ***Dirofilaria immitis*** | qPCR | ***cox1*** | Primer | S0624_F | TAGAGGGTCAGCCTGAGTTATC | (Panetta et al., 2021) |
|  |  |  |  | S0626_R | AGTAGAACGTATATTCTGAACAGTAACC |  |
|  |  |  | Probe | S0625_P | AGAACCAATACCAACAGTATGAAGACC |  |
| ***Dirofilaria immitis*-associated *Wolbachia*** | qPCR | ***ftsZ*** | Primer | WDiro.ftsZ.490-F | AAGCCATTTRGCTTYGAAGGTG | (Laidoudi et al., 2020) |
|  |  |  |  | WDiro.ftsZ.600-R | AAACAAGTTTTGRTTTGGAATAACAAT |  |
|  |  |  | Probe | WDimm.ftsZ.523-P | CGTATTGCAGAGCTCGGATTA |  |
| ***Dirofilaria immitis* L42411 SNP** | Illumina amplicon NGS | **L42411** | Primer with Illumina overhang adapter | S1020_NODE_42411_FOR | TCGTCGGCAGCGTCAGATGTGTATAAGAGACAGTTCTATCGAAAACCTTCCAG | (Bourguinat et al., 2015) |
|  |  |  |  | S1021_NODE_42411_REV | GTCTCGTGGGCTCGGAGATGTGTATAAGAGACAGAGGTTGCAAAAGTTGCAATG |  |
| ***Dirofilaria immitis* L21554 SNP** | Illumina amplicon NGS | **L21554** | Primer with Illumina overhang adapter | S1012_NODE_21554_FOR | TCGTCGGCAGCGTCAGATGTGTATAAGAGACAGCATCGTTGTCAACTTCCTGC | (Bourguinat et al., 2015) |
|  |  |  |  | S1013_NODE_21554_REV | GTCTCGTGGGCTCGGAGATGTGTATAAGAGACAGGAAATTTGAAAATGGGTACT |  |
| ***Dirofilaria immitis* L45689 SNP** | Illumina amplicon NGS | **L45689** | Primer with Illumina overhang adapter | S1022_NODE_45689_FOR | TCGTCGGCAGCGTCAGATGTGTATAAGAGACAGACGCAGGAAAGCTTTAATGG | (Bourguinat et al., 2015) |
|  |  |  |  | S1023_NODE_45689_REV | GTCTCGTGGGCTCGGAGATGTGTATAAGAGACAGATCATCATTTTATCAATTCC |  |
| ***Dirofilaria immitis* L9400 SNP** | Illumina amplicon NGS | **L9400** | Primer with Illumina overhang adapter | S1026_NODE_9400_FOR | TCGTCGGCAGCGTCAGATGTGTATAAGAGACAGGTTATTTGCACTACTCTCCC | (Bourguinat et al., 2015) |
|  |  |  |  | S1027_NODE_9400_REV | GTCTCGTGGGCTCGGAGATGTGTATAAGAGACAGTGGCGTACTGATCACATTGG |  |
| ***Dirofilaria immitis* L20587 SNP** | Illumina amplicon NGS | **L20587** | Primer with Illumina overhang adapter | S1010_NODE_20587_FOR | TCGTCGGCAGCGTCAGATGTGTATAAGAGACAGTCGATCATTTAGTAACAACG | (Bourguinat et al., 2015) |
|  |  |  |  | S1011_NODE_20587_REV | GTCTCGTGGGCTCGGAGATGTGTATAAGAGACAGTTGCGTTACAGCGCCAAATC |  |
| ***Dirofilaria immitis* L15709A SNP** | Illumina amplicon NGS | **L15709A** | Primer with Illumina overhang adapter | S1008_NODE_15709_A_FOR | TCGTCGGCAGCGTCAGATGTGTATAAGAGACAGGGCCAATAAATAAAGGCTA | (Bourguinat et al., 2015) |
|  |  |  |  | S1009_NODE_15709_A_REV | GTCTCGTGGGCTCGGAGATGTGTATAAGAGACAGGTTTTCTGGAATTATCAGAC |  |
| ***Dirofilaria immitis* L30575 SNP** | Illumina amplicon NGS | **L30575** | Primer with Illumina overhang adapter | S1018_NODE_30575_FOR | TCGTCGGCAGCGTCAGATGTGTATAAGAGACAGCGAGGTAAAGCACACAGAAG | (Bourguinat et al., 2015) |
|  |  |  |  | S1019_NODE_30575_REV | GTCTCGTGGGCTCGGAGATGTGTATAAGAGACAGCAACAAAATGCCGCAGATGG |  |
