## Supplementary Table S2 for "Exploration of the sensitivity to macrocyclic lactones in the canine heartworm (*Dirofilaria immitis*) in Australia using phenotypic and genotypic approaches": MS_RoseHW_Supplementary_Table S2.docx

**Supplementary Table S2:** SNPs in this study adopted from previous 5-SNPs models from the USA

| **SNP position** | **Node / locus** | **SNP included in 5-SNPs model** | | **SNP included** |
| --- | --- | --- | --- | --- |
|  |  | (Ballesteros et al., 2018) | (Bourguinat et al., 2017) | **This study** |
| nDi.2.2.scaf00046:76278 | 15709_A | Yes | No | Yes |
| nDi.2.2.scaf00046:22857 | 30575 | Yes | No | Yes |
| nDi.2.2.scaf00046:222254 | 21554 | Yes | Yes | Yes |
| nDi.2.2.scaf00185:10639 | 45689 | Yes | Yes | Yes |
| nDi.2.2.scaf00185:62174 | 20587 | Yes | Yes | Yes |
| nDi.2.2.scaf00001-466197 | 42411 | No | Yes | Yes |
| nDi.2.2.scaf00005-662854 | 9400 | No | Yes | Yes |
